# Major histocompatibility complex haplotyping and long-amplicon allele discovery in cynomolgus macaques from Chinese breeding facilities

**DOI:** 10.1101/084947

**Authors:** Julie A. Karl, Michael E. Graham, Roger W. Wiseman, Katelyn E. Heimbruch, Samantha M. Gieger, Gaby G. M. Doxiadis, Ronald E. Bontrop, David H. O’Connor

**Author notes:** Corresponding Author: David H. O’Connor, Department of Pathology and Laboratory Medicine, University of Wisconsin-Madison, 585 Science Drive, Madison, WI, 53711.

## Abstract

Very little is currently known about the major histocompatibility complex (MHC) region of cynomolgus macaques (*Macaca fascicularis*; *Mafa*) from Chinese breeding centers. We performed comprehensive MHC class I haplotype analysis of 100 cynomolgus macaques from two different centers, with animals from different reported original geographic origins (Vietnamese, Cambodian, and Cambodian/Indonesian mixed-origin). Many of the samples were of known relation to each other (sire, dam, and progeny sets), making it possible to characterize lineage-level haplotypes in these animals. We identified 52 *Mafa-A* and 74 *Mafa-B* haplotypes in this cohort, many of which were restricted to specific sample origins. We also characterized full-length MHC class I transcripts using Pacific Biosciences (PacBio) RS II single-molecule real-time (SMRT) sequencing. This technology allows for complete read-through of unfragmented MHC class I transcripts (~1,100 bp in length), so no assembly is required to unambiguously resolve novel full-length sequences. Overall, we identified 313 total full-length transcripts in a subset of 72 cynomolgus macaques from these Chinese breeding facilities; 131 of these sequences were novel and an additional 116 extended existing short database sequences to span the complete open reading frame. This significantly expands the number of *Mafa-A*, *Mafa-B*, and *Mafa-I* full-length alleles in the official cynomolgus macaque MHC class I database. The PacBio technique described here represents a general method for full-length allele discovery and genotyping that can be extended to other complex immune loci such as MHC class II, killer immunoglobulin-like receptors, and Fc gamma receptors.

## INTRODUCTION

Cynomolgus macaques (*Macaca fascicularis*; *Mafa*) are native to much of continental and insular Southeast Asia, but there are no feral cynomolgus macaques living in China. However, Chinese breeding facilities are one of the largest exporters of cynomolgus macaques for use in biomedical research. According to the Centers for Disease Control and Prevention, China provided over 70% of the total cynomolgus macaques imported into the United States in fiscal year 2014. These animals are derived from other geographic populations of cynomolgus macaques, most likely from Thailand, Laos, Vietnam, Cambodia, and Malaysia, but also possibly including animals from Indonesia and the Philippines. Cynomolgus macaques from Chinese breeding facilities therefore are genetically heterogeneous, and this heterogeneity may be particularly biologically relevant in highly diverse genomic regions like the major histocompatibility complex (MHC).

The MHC is both highly complex and important – the gene products encoded by the MHC present peptides to T-cells, playing an essential role in immune response to pathogens and other non-self peptides (Bontrop 2006). In macaques, the MHC class I region is highly polymorphic and extensively duplicated (Daza-Vamenta et al. 2004; Otting et al. 2005), with different chromosomes carrying different numbers of functional *Mafa-A* and *Mafa-B* genes. Many investigators have examined the MHC class I region of cynomolgus macaques from various origins (Budde et al. 2010; Campbell et al. 2008; Kita et al. 2009; Lawrence et al. 2012; Ling et al. 2012; Otting et al. 2007, 2009, 2012; Pendley et al. 2008; Saito et al. 2012; Shiina et al. 2015; Uda et al. 2004, 2005; Zhang et al. 2012; Zhuo et al. 2011), but only a limited number of studies have looked at cynomolgus macaques from Chinese breeding facilities (Krebs et al. 2005; Westbrook et al. 2015).

Here we describe a comprehensive study of 100 samples collected from two different breeding facilities in China that include individuals reported to be of Vietnamese, Cambodian, and mixed Cambodian/Indonesian origins. Many of the samples were related to at least one other animal in the study, allowing us to characterize lineage-level haplotypes based on shared Illumina short-read transcript sequences. We then performed full-length MHC class I transcript analysis on a subset of the samples using the PacBio long-read single-molecule real-time (SMRT) sequencing system. This analysis improved on a previous MHC class I study using PacBio circular consensus SMRT sequencing (Westbrook et al. 2015) by using more advanced sequencing chemistry, and a data analysis pipeline focused on the highest quality reads that did not need to account for sequencing reads with insertion/deletion errors.

## MATERIALS AND METHODS

### Animals

A total of 100 whole blood samples were collected from two different breeding facilities in China. Thirty samples from reported Vietnamese-origin animals were obtained from Yulin Hongfeng Experimental Animal’s Domesticating and Breeding Center (Guangxi, China). A total of 70 samples were obtained from Hainan Jingang Laboratory Animal Co., Ltd. (Hainan, China) - 57 samples were obtained from reported Cambodian-origin animals, and 13 samples were from mixed Cambodian/Indonesian-origin animals. Wherever possible, sire/dam/progeny trio sets or parent/progeny duos were collected to obtain samples that share MHC haplotypes that are identical by descent; a total of 71 samples across both facilities were directly related to at least one other animal in the cohort. Samples were collected into TRIzol Reagent (ThermoFisher Scientific, Waltham, MA, USA) to stabilize the total RNA for shipment from China.

### RNA isolation, cDNA synthesis, and PCR preparation

Total RNA was isolated from the blood samples and cDNA was produced from the isolated RNA using the RevertAid First Strand cDNA Synthesis Kit (ThermoFisher Scientific) following manufacturer’s protocols. For Illumina MiSeq genotyping, a 195 bp amplicon spanning the most polymorphic region of MHC class I molecules was generated using primers designed in conserved regions of exon 2, as shown in Supplemental Fig. 1. Amplification was performed using a 4-primer system to add unique barcodes to each sample, using consensus sequence adapters and barcodes designed by the Fluidigm Corporation (San Francisco, CA, USA), as detailed in Supplemental File 1.

**Fig. 1.**
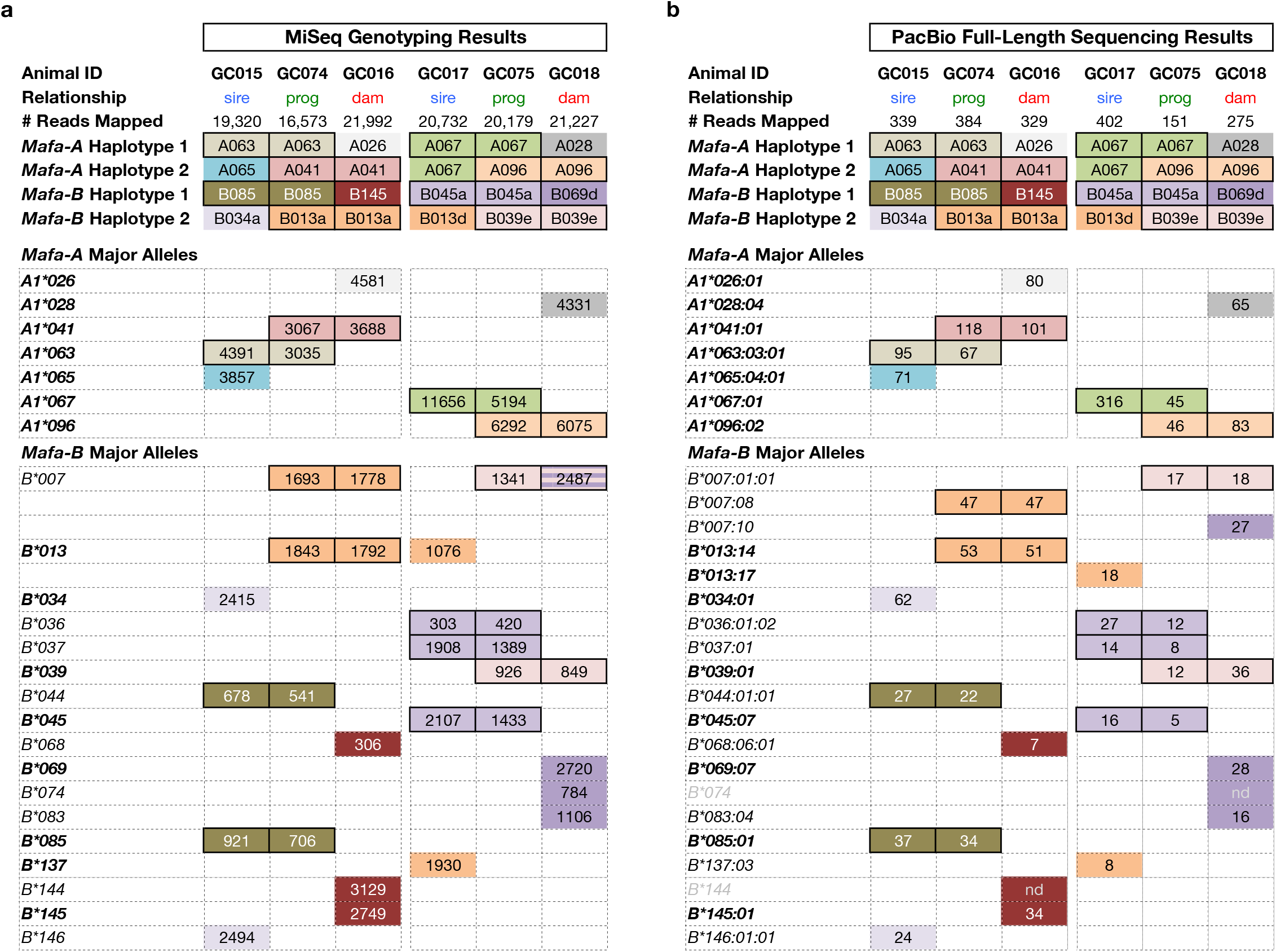
Representative trio set results by a) MiSeq genotyping and b) PacBio full-length sequencing methods. The major transcripts for two representative trio sets are shown here, with progeny located between the sire and dam, showing direct transcript inheritance from parents to offspring. Inherited haplotypes are outlined in solid boxes. Groups of *Mafa-A* and *Mafa-B* transcripts co-inherited together as a haplotype are colored correspondingly. Transcripts shown in bold font are diagnostic major sequences used to name the haplotypes. Transcripts shown in grey font were only observed by MiSeq. Values indicate the number of reads for each sequence per sample

For PacBio RS II full-length MHC class I transcript sequencing of 72 samples from the full cohort, an ~1,150 bp amplicon with primers located in the 5‘UTR and 3’UTR of MHC class I sequences was generated as shown in Supplemental Fig. 2. A cocktail of two forward primers and three reverse primers was used to maximize homology with the majority of MHC class I sequences. A unique set of barcoded primers was used for each sample, with the barcodes integrated onto the ends of the sequence-specific oligos at synthesis. Barcodes are available from Pacific Biosciences (Menlo Park, CA, USA). Primer sequences are shown in Supplemental Fig. 2, and PCR conditions are detailed in Supplemental File 1.

**Fig. 2.**
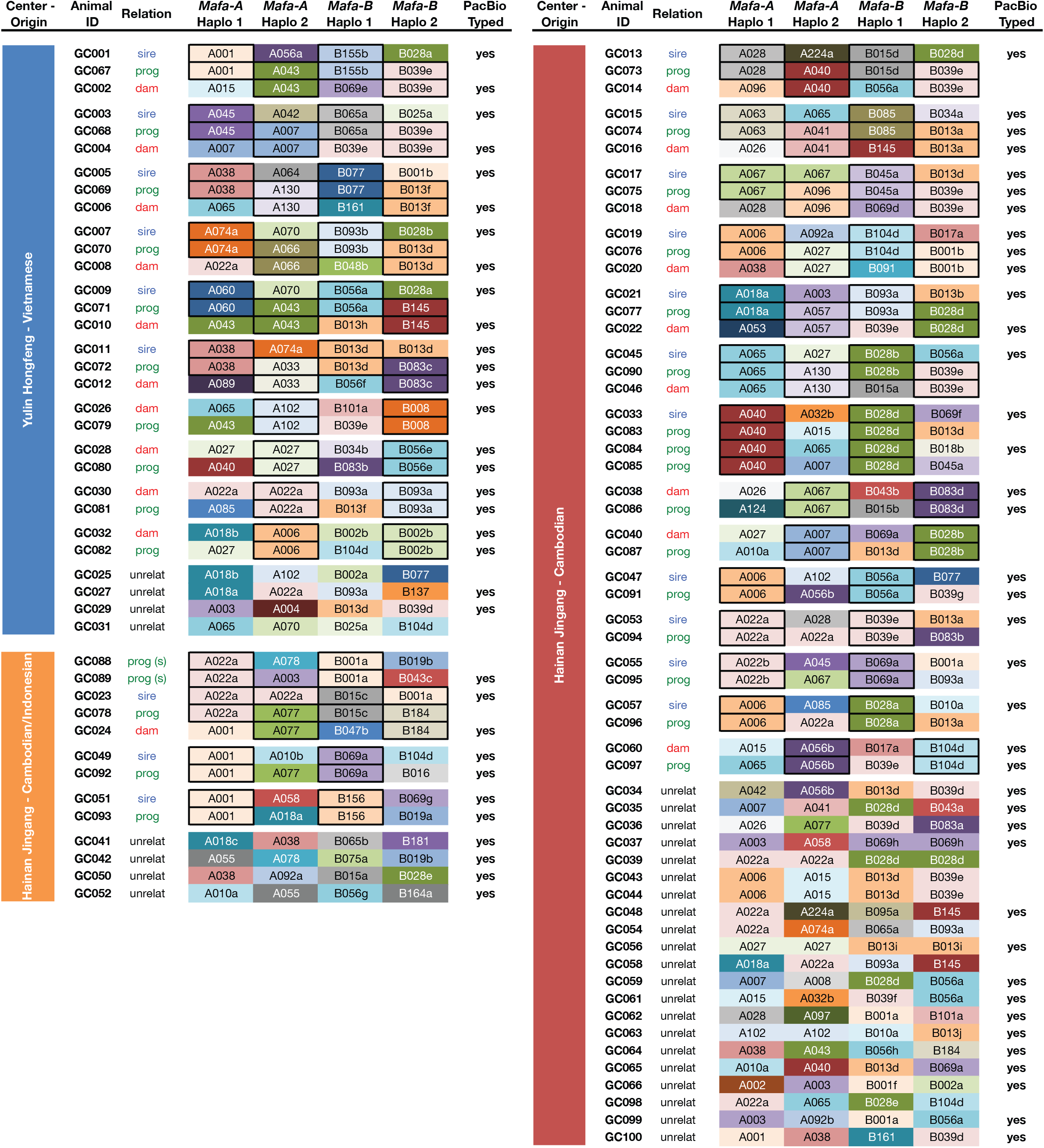
Summary of MHC class I *Mafa-A* and *Mafa-B* haplotypes observed in each macaque. Samples are organized by geographic origin into family groups, and directly inherited haplotypes are outlined with solid boxes. The full list of transcripts associated with each haplotype are available in Supplemental Table 1

**Table 1:** Full-length transcripts observed by PacBio sequencing

| Allele Name | Full-Length Accession Number | # Animals | # Reads | ORF Length (bp) | Category | Longest Previous Length (bp) | Previous Accession Number(s) |
| --- | --- | --- | --- | --- | --- | --- | --- |
| *Mafa-A1\*001:01:02* | LN851916 | 3 | 123 | 1098 | extension | 1062 | FR682846 |
| *Mafa-A1\*001:01:03* | LN994391 | 1 | 31 | 1098 | novel | - | - |
| *Mafa-A1\*001:02:01* | LN998188 | 1 | 54 | 1098 | extension | 1039 | AY958101 |
| *Mafa-A1\*001:05* | LN851917 | 2 | 51 | 1098 | novel | - | - |
| *Mafa-A1\*002:02* | LN994392 | 1 | 35 | 1098 | extension | 1003 | AB447609 |
| *Mafa-A1\*003:01* | LN994393 | 1 | 48 | 1098 | extension | 1065 | AM295829 |
| *Mafa-A1\*003:03* | LN994394 | 1 | 78 | 1098 | extension | 1062 | FR682835, HQ113982, HQ131743 |
| *Mafa-A1\*003:05* | LN994395 | 1 | 73 | 1098 | extension | 544 | HQ131758 |
| *Mafa-A1\*003:06* | LN851918 | 3 | 337 | 1098 | novel | - | - |
| *Mafa-A1\*003:07* | LN994396 | 1 | 53 | 1098 | novel | - | - |
| *Mafa-A1\*004:02* | LN994397 | 1 | 60 | 1098 | extension | 1003 | AB448751, HQ113954, HQ131752 |
| *Mafa-A1\*006:01:01* | LN851919 | 7 | 528 | 1098 | extension | 1062 | AY958098, FR682819 |
| *Mafa-A1\*007:01* | LN851920 | 2 | 627 | 1098 | extension | 1042 | AY958089, HQ113975 |
| *Mafa-A1\*007:06* | LN994398 | 1 | 163 | 1098 | extension | 1065 | FR691532, KP307886 |
| *Mafa-A1\*008:01* | KP307887 | 1 | 178 | 1098 | extant | 1098 | AY958104, HQ113960, HQ131766, KP307887 |
| *Mafa-A1\*010:04* | EU203706 | 1 | 117 | 1098 | extant | 1098 | EU203706 |
| *Mafa-A1\*010:05* | EU203707 | 1 | 115 | 1098 | extant | 1098 | EU203707, FR686840 |
| *Mafa-A1\*010:07* | LN851921 | 1 | 50 | 1098 | novel | - | - |
| *Mafa-A1\*015:01* | LN851922 | 3 | 309 | 1098 | extension | 1039 | AY958087 |
| *Mafa-A1\*018:01* | LN851923 | 2 | 337 | 1098 | extension | 1062 | AY958090, FR682839 |
| *Mafa-A1\*018:06* | FM246489 | 2 | 279 | 1098 | extant | 1098 | FM246489 |
| *Mafa-A1\*018:08* | LN994399 | 1 | 36 | 1098 | novel | - | - |
| *Mafa-A1\*022:01* | LN994400 | 1 | 17 | 1098 | extension | 1042 | AY958096, LN994401 |
| *Mafa-A1\*022:04* | LN851924 | 5 | 634 | 1098 | extension | 822 | HM236273, HQ113976, HQ131745 |
| *Mafa-A1\*022:05* | LN851925 | 1 | 141 | 1098 | extension | 1062 | FR682826, HQ113977, HQ131770 |
| *Mafa-A1\*022:06* | KY047492 | 1 | 293 | 1098 | extension | 1062 | FR682828, HQ131736 |
| *Mafa-A1\*022:09:01* | KP307910 | 1 | 58 | 1098 | extant | 1098 | KP307910 |
| *Mafa-A1\*022:09:02* | LN851926 | 1 | 47 | 1098 | novel | - | - |
| *Mafa-A1\*026:01* | KP307888 | 3 | 306 | 1098 | extant | 1098 | AY958119, HQ131773, KP307888 |
| *Mafa-A1\*027:01* | LN851927 | 6 | 705 | 1098 | extension | 1065 | AY958103, KP307889 |
| *Mafa-A1\*027:02* | LN994402 | 1 | 41 | 1098 | extension | 1042 | AY958113 |
| *Mafa-A1\*027:03* | LN994403 | 1 | 102 | 1098 | extension | 1003 | AB447615 |
| *Mafa-A1\*028:01* | LN851928 | 1 | 57 | 1098 | extension | 1062 | FR682824, HQ113969, HQ131769 |
| *Mafa-A1\*028:03* | LN994404 | 1 | 225 | 1098 | novel | - | - |
| *Mafa-A1\*028:04* | LN994405 | 1 | 65 | 1098 | novel | - | - |
| *Mafa-A1\*028:05* | LN994406 | 1 | 22 | 1098 | novel | - | - |
| *Mafa-A1\*032:03:02* | LN851929 | 2 | 195 | 1098 | extension | 1062 | FR682840 |
| *Mafa-A1\*033:02* | LN851930 | 2 | 157 | 1098 | novel | - | - |
| *Mafa-A1\*038:01* | LN851931 | 8 | 894 | 1098 | extension | 1065 | AM295825, AY958114, FR682847, HQ131737 |
| *Mafa-A1\*040:01:01* | LN851932 | 3 | 275 | 1089 | extension | 1062 | AY958115, FR682842 |
| *Mafa-A1\*040:03* | LN994407 | 1 | 70 | 1089 | extension | 1042 | AY958095 |
| *Mafa-A1\*041:01* | LN851933 | 3 | 350 | 1098 | extension | 1042 | AY958116 |
| *Mafa-A1\*042:01* | LN994408 | 1 | 109 | 1098 | extension | 1039 | AY958117 |
| *Mafa-A1\*042:02* | LN851934 | 1 | 85 | 1098 | extension | 1062 | FR682858 |
| *Mafa-A1\*043:01* | LN994409 | 1 | 144 | 1098 | extension | 1062 | AY958118, FR686841 |
| *Mafa-A1\*043:10* | KP307853 | 2 | 127 | 1098 | extant | 1098 | KP307853, KP307884 |
| *Mafa-A1\*043:11* | LN994410 | 1 | 43 | 1098 | novel | - | - |
| *Mafa-A1\*045:01* | LN851935 | 1 | 107 | 1098 | extension | 1062 | AY958088, AY958120, FR682843 |
| *Mafa-A1\*045:03* | LN994411 | 1 | 114 | 1098 | novel | - | - |
| *Mafa-A1\*053:01* | LN994412 | 1 | 57 | 1098 | extension | 1065 | AB447562, AM295839, HQ113973, HQ131754 |
| *Mafa-A1\*055:01* | LN994413 | 1 | 108 | 1098 | extension | 1065 | AM295840 |
| *Mafa-A1\*055:02* | LN994414 | 1 | 214 | 1098 | novel | - | - |
| *Mafa-A1\*056:03* | KP307890 | 5 | 428 | 1098 | extant | 1098 | KP307890 |
| *Mafa-A1\*057:02* | LN994415 | 1 | 85 | 1098 | extension | 1065 | KP307880 |
| *Mafa-A1\*058:02* | LN851936 | 2 | 194 | 1098 | extension | 1062 | FR682836 |
| *Mafa-A1\*060:04* | LN851937 | 1 | 51 | 1098 | extension | 1003 | AB447574 |
| *Mafa-A1\*063:03:01* | LN851938 | 2 | 162 | 1098 | extension | 1062 | AY958099, FR682832 |
| *Mafa-A1\*064:03* | LN994416 | 1 | 287 | 1098 | extension | 1003 | AB447608 |
| *Mafa-A1\*065:02* | LN851939 | 1 | 36 | 1098 | extension | 1003 | AB447611 |
| *Mafa-A1\*065:03* | LN851940 | 3 | 144 | 1098 | extension | 1062 | AB447614, FR682853 |
| *Mafa-A1\*065:04:01* | LN851941 | 1 | 71 | 1098 | extension | 1062 | AB448748, FR682825, FR682850 |
| *Mafa-A1\*065:05* | LN994417 | 1 | 59 | 1098 | novel | - | - |
| *Mafa-A1\*066:06* | LN851942 | 1 | 102 | 1098 | extension | 721 | HQ113962, HQ131765 |
| *Mafa-A1\*067:01* | KP307908 | 4 | 584 | 1098 | extant | 1098 | AM295855, FR682827, KP307908 |
| *Mafa-A1\*070:01* | EU203708 | 2 | 74 | 1098 | extant | 1098 | AM295858, EU203708, FR682831 |
| *Mafa-A1\*074:01:01* | LN851943 | 2 | 220 | 1098 | extension | 1003 | AB447596 |
| *Mafa-A1\*077:01:02* | LN994418 | 1 | 81 | 1098 | extension | 1003 | AB447613 |
| *Mafa-A1\*077:02:01* | LN994419 | 1 | 101 | 1098 | novel | - | - |
| *Mafa-A1\*077:02:02* | LN994420 | 1 | 275 | 1098 | novel | - | - |
| *Mafa-A1\*078:01:01* | AB154767 | 1 | 111 | 1098 | extant | 1098 | AB154767 |
| *Mafa-A1\*085:02* | LN851944 | 2 | 198 | 1098 | extension | 1062 | FR682820 |
| *Mafa-A1\*086:01:02* | LN851945 | 2 | 12 | 981 | novel | - | - |
| *Mafa-A1\*086:03* | LN994421 | 1 | 5 | 981 | extension | 822 | GU130454 |
| *Mafa-A1\*086:05* | LN994422 | 1 | 5 | 981 | novel | - | - |
| *Mafa-A1\*089:06* | LN851946 | 1 | 68 | 1098 | extension | 1003 | AB447610 |
| *Mafa-A1\*090:02:02* | LN994423 | 1 | 10 | 1098 | novel | - | - |
| *Mafa-A1\*090:03* | LN851947 | 4 | 19 | 1098 | novel | - | - |
| *Mafa-A1\*092:01* | LN994424 | 1 | 41 | 1098 | extension | 1062 | AB447594, FR682823, FR682833, HQ113961, HQ131768 |
| *Mafa-A1\*092:03* | LN994425 | 1 | 123 | 1098 | extension | 1065 | FM246488 |
| *Mafa-A1\*092:04* | KP307869 | 1 | 103 | 1098 | extant | 1098 | KP307869 |
| *Mafa-A1\*095:02* | LN994426 | 1 | 99 | 1098 | novel | - | - |
| *Mafa-A1\*096:02* | LN994427 | 2 | 129 | 1098 | novel | - | - |
| *Mafa-A1\*097:02* | LN994428 | 1 | 119 | 1098 | novel | - | - |
| *Mafa-A1\*101:02:02* | LN851948 | 1 | 67 | 1098 | novel | - | - |
| *Mafa-A1\*102:03:01* | LN851949 | 3 | 256 | 1098 | novel | - | - |
| *Mafa-A1\*102:03:02* | LN994429 | 1 | 146 | 1098 | novel | - | - |
| *Mafa-A1\*124:01* | LN994430 | 1 | 63 | 1098 | extension | 641 | AB583237 |
| *Mafa-A1\*130:01* | LN851950 | 1 | 66 | 1098 | extension | 1080 | KP307844 |
| *Mafa-A2\*01:01* | LN994431 | 1 | 62 | 1098 | extension | 1062 | FR682848, HQ113963, HQ131750 |
| *Mafa-A2\*05:06:02* | LN994432 | 4 | 6 | 1098 | extension | 822 | EF550522 |
| *Mafa-A2\*05:06:04* | LN994438 | 2 | 14 | 1098 | novel | - | - |
| *Mafa-A2\*05:22* | LN851952 | 2 | 14 | 1101 | extension | 822 | EF589357 |
| *Mafa-A2\*05:25:01* | LN994433 | 5 | 16 | 1098 | extension | 822 | EF589360, HP823075 |
| *Mafa-A2\*05:31* | LN851953 | 2 | 18 | 1101 | extension | 822 | EF550520 |
| *Mafa-A2\*05:40* | LN994434 | 2 | 5 | 1098 | extension | 822 | GQ131777 |
| *Mafa-A2\*05:42* | LN994435 | 1 | 20 | 1083 | extension | 822 | GQ131775 |
| *Mafa-A2\*05:45* | LN994436 | 1 | 4 | 1098 | extension | 822 | GQ131771 |
| *Mafa-A2\*05:46* | LN851954 | 2 | 9 | 1101 | extension | 1065 | GQ131770, HQ230581 |
| *Mafa-A2\*05:50* | GU063713 | 1 | 1 | 1098 | extant | 1098 | GU063713, LC053847 |
| *Mafa-A2\*05:51* | LN851955 | 1 | 3 | 1101 | extension | 822 | GU130445 |
| *Mafa-A2\*05:55* | LN851956 | 5 | 48 | 1101 | extension | 1080 | KP307855 |
| *Mafa-A2\*05:57:01* | LN851951 | 6 | 34 | 1098 | extension | 822 | HQ992789 |
| *Mafa-A2\*05:58* | LN994437 | 1 | 5 | 1098 | novel | - | - |
| *Mafa-A2\*05:59* | LN994439 | 2 | 6 | 1098 | novel | - | - |
| *Mafa-A2\*24:07* | LN851957 | 1 | 49 | 1098 | novel | - | - |
| *Mafa-A2\*24:08* | LN994440 | 1 | 211 | 1098 | novel | - | - |
| *Mafa-A3\*13:02* | LN851958 | 2 | 27 | 1098 | extension | 1042 | AY958111 |
| *Mafa-A3\*13:07* | LN851959 | 4 | 55 | 1098 | extension | 1062 | FR682834, GQ131768, HM161349, HQ113968, HQ131739 |
| *Mafa-A3\*13:14N* | LN851960 | 2 | 5 | 1011 | extension | 822 | GU130464 |
| *Mafa-A3\*13:15* | LN994441 | 1 | 6 | 1098 | extension | 641 | AB583238 |
| *Mafa-A3\*13:17:02* | LN851961 | 1 | 5 | 1098 | novel | - | - |
| *Mafa-A3\*13:20* | LN851962 | 1 | 15 | 1098 | novel | - | - |
| *Mafa-A3\*13:21* | LN994442 | 1 | 6 | 1098 | novel | - | - |
| *Mafa-A3\*13:22* | LN994443 | 1 | 14 | 1098 | novel | - | - |
| *Mafa-A4\*01:02* | LN851963 | 2 | 9 | 1098 | extension | 1042 | AY958121 |
| *Mafa-A4\*01:08* | LN994444 | 6 | 11 | 1011 | extension | 822 | GQ131766 |
| *Mafa-A4\*01:12* | LN994445 | 1 | 6 | 1011 | novel | - | - |
| *Mafa-A4\*14:01* | LN851964 | 9 | 31 | 1011 | extension | 1065 | AM295880 |
| *Mafa-A4\*14:03* | LN851965 | 21 | 110 | 1101 | extension | 1065 | FR682829, GQ131781, HQ113965, HQ131747, LC043320 |
| *Mafa-A4\*14:04* | LN851967 | 1 | 3 | 1101 | extension | 822 | GQ131767 |
| *Mafa-A4\*14:13* | KP307856 | 1 | 22 | 1092 | extant | 1092 | KP307856 |
| *Mafa-A4\*14:15* | LN851968 | 1 | 6 | 1107 | novel | - | - |
| *Mafa-A4\*14:16* | LN851966 | 1 | 3 | 1101 | novel | - | - |
| *Mafa-A4\*14:18* | LN994446 | 1 | 4 | 1107 | novel | - | - |
| *Mafa-A4\*14:19* | LN994447 | 1 | 7 | 1101 | novel | - | - |
| *Mafa-A5\*30:06* | KP307871 | 1 | 1 | 1089 | extant | 1089 | KP307871 |
| *Mafa-B\*001:01:01* | KP307893 | 7 | 239 | 1089 | extant | 1089 | AY958126, KP307893 |
| *Mafa-B\*001:01:02* | LN994449 | 1 | 54 | 1089 | novel | - | - |
| *Mafa-B\*001:02* | LN994448 | 1 | 19 | 1089 | novel | - | - |
| *Mafa-B\*002:04* | LN994450 | 3 | 472 | 1089 | extension | 822 | HM161352, HQ114001, HQ131729, HQ131749 |
| *Mafa-B\*006:01:02* | LN994451 | 1 | 12 | 1089 | novel | - | - |
| *Mafa-B\*007:01:01* | GU063753 | 17 | 606 | 1089 | extant | 1089 | AY958137, GU063753, HM161384, HM236279 |
| *Mafa-B\*007:01:05* | LN851969 | 6 | 155 | 1089 | extension | 822 | HM161391, HQ114005, HQ131716 |
| *Mafa-B\*007:05* | LN851970 | 4 | 98 | 1089 | extension | 544 | HQ131707 |
| *Mafa-B\*007:06:02* | LN994453 | 1 | 48 | 1089 | novel | - | - |
| *Mafa-B\*007:07* | LN851971 | 3 | 53 | 1089 | novel | - | - |
| *Mafa-B\*007:08* | LN851972 | 7 | 235 | 1089 | novel | - | - |
| *Mafa-B\*007:10* | LN994452 | 1 | 27 | 1089 | novel | - | - |
| *Mafa-B\*008:01* | LN994454 | 1 | 56 | 1089 | extension | 544 | HQ131700 |
| *Mafa-B\*010:01* | AB195452 | 2 | 112 | 1089 | extant | 1089 | AB195452 |
| *Mafa-B\*011:01* | AY958143 | 1 | 42 | 1089 | extant | 1089 | AY958143, EU203717, GU063737 |
| *Mafa-B\*013:03* | LN994455 | 2 | 45 | 1089 | extension | 822 | FJ178813 |
| *Mafa-B\*013:06* | LN851973 | 1 | 32 | 1089 | extension | 822 | GQ131788, HQ113998, HQ131703 |
| *Mafa-B\*013:09* | KP307894 | 3 | 269 | 1089 | extant | 1089 | FJ178814, HQ131706, KP307894 |
| *Mafa-B\*013:10* | LN851974 | 2 | 42 | 1089 | extension | 1056 | HM161366, HQ131722, KP307895 |
| *Mafa-B\*013:11* | LN994456 | 1 | 45 | 1089 | extension | 1056 | FM212794 |
| *Mafa-B\*013:13* | LN851975 | 1 | 27 | 1089 | novel | - | - |
| *Mafa-B\*013:14* | LN851976 | 2 | 104 | 1089 | novel | - | - |
| *Mafa-B\*013:15* | LN994457 | 1 | 44 | 1089 | novel | - | - |
| *Mafa-B\*013:16* | LN994458 | 1 | 118 | 1089 | novel | - | - |
| *Mafa-B\*013:17* | LN994459 | 1 | 18 | 1089 | novel | - | - |
| *Mafa-B\*013:18* | LN994460 | 1 | 4 | 1089 | novel | - | - |
| *Mafa-B\*014:01* | LN851977 | 2 | 14 | 1089 | extension | 1056 | FM212831 |
| *Mafa-B\*014:02* | LN851978 | 1 | 8 | 1089 | novel | - | - |
| *Mafa-B\*015:01* | LN851979 | 2 | 49 | 1089 | extension | 1042 | AY958132, HM235701 |
| *Mafa-B\*015:05* | LN994461 | 1 | 42 | 1089 | novel | - | - |
| *Mafa-B\*015:06* | LN994462 | 1 | 106 | 1089 | novel | - | - |
| *Mafa-B\*016:01* | EU203701 | 1 | 118 | 1092 | extant | 1092 | EU203701 |
| *Mafa-B\*016:02* | LN994463 | 1 | 61 | 1092 | novel | - | - |
| *Mafa-B\*017:02* | KP307896 | 2 | 43 | 1080 | extant | 1080 | HQ131701, KP307896 |
| *Mafa-B\*018:01:01* | KP307897 | 3 | 178 | 1089 | extant | 1089 | AY958138, KP307897 |
| *Mafa-B\*018:03* | LN994464 | 1 | 13 | 1089 | novel | - | - |
| *Mafa-B\*019:03* | DQ979879 | 1 | 7 | 1086 | extant | 1086 | DQ979879, GU063743 |
| *Mafa-B\*019:04* | LN994465 | 1 | 33 | 1086 | novel | - | - |
| *Mafa-B\*021:02* | KP307859 | 8 | 127 | 1077 | extant | 1077 | KP307859, KP307845 |
| *Mafa-B\*021:04* | LN994466 | 1 | 10 | 1077 | novel | - | - |
| *Mafa-B\*021:05* | LN994467 | 2 | 37 | 1077 | novel | - | - |
| *Mafa-B\*025:02* | LN851980 | 1 | 110 | 1086 | novel | - | - |
| *Mafa-B\*027:03:01* | LN851982 | 1 | 3 | 1096+ | extension | 659 | HG977466 |
| *Mafa-B\*027:03:02* | LN851981 | 2 | 19 | 1096+ | novel | - | - |
| *Mafa-B\*028:02* | KP307898 | 6 | 18 | 1089 | extant | 1089 | AY958130, KP307898 |
| *Mafa-B\*028:03* | LN994468 | 1 | 6 | 1089 | extension | 1042 | AY958128, LC043325 |
| *Mafa-B\*028:04* | EU046324 | 2 | 20 | 1089 | extant | 1089 | EU046324, HM235716, HQ131708 |
| *Mafa-B\*030:01:01* | LN851983 | 7 | 277 | 1089 | extension | 1042 | AY958133 |
| *Mafa-B\*030:01:05* | LN994472 | 2 | 22 | 1089 | novel | - | - |
| *Mafa-B\*030:02* | LN994469 | 1 | 16 | 1089 | extension | 1042 | AY958134 |
| *Mafa-B\*030:03:01* | KP307900 | 2 | 10 | 1089 | extant | 1089 | AY958135, HQ131687, KP307900 |
| *Mafa-B\*030:03:02* | LN994470 | 5 | 18 | 1089 | extension | 822 | FJ719167, HQ131726 |
| *Mafa-B\*030:04* | LN994471 | 1 | 48 | 1089 | extension | 822 | FJ719169 |
| *Mafa-B\*030:15* | KP307857 | 2 | 34 | 1089 | extant | 1089 | KP307857 |
| *Mafa-B\*030:16* | LN851984 | 2 | 22 | 1089 | novel | - | - |
| *Mafa-B\*030:17* | KP307899 | 5 | 28 | 1089 | extant | 1089 | FJ178822, KP307899 |
| *Mafa-B\*030:18* | LN994473 | 2 | 5 | 1089 | novel | - | - |
| *Mafa-B\*030:19* | LN994474 | 1 | 4 | 1089 | novel | - | - |
| *Mafa-B\*031:03* | LN994475 | 2 | 21 | 1095 | novel | - | - |
| *Mafa-B\*034:01* | LN851985 | 1 | 62 | 1089 | extension | 1042 | AY958127 |
| *Mafa-B\*034:03* | LN851986 | 2 | 131 | 1089 | extension | 819 | HQ113997, HQ131704 |
| *Mafa-B\*034:04* | LN994476 | 1 | 34 | 1089 | extension | 1056 | KP307879 |
| *Mafa-B\*036:01:02* | EU606046 | 2 | 39 | 1080 | extant | 1080 | EU606046, FM212796 |
| *Mafa-B\*037:01* | AY958149 | 2 | 22 | 1089 | extant | 1089 | AY958149, HQ114004, HQ131697 |
| *Mafa-B\*038:01* | KP307860 | 2 | 31 | 1089 | extant | 1089 | KP307860 |
| *Mafa-B\*039:01* | KP307902 | 11 | 674 | 1089 | extant | 1089 | HM161344, HQ113949, HQ113992, HQ114010, HQ131713, HQ131762, KP307902 |
| *Mafa-B\*039:02* | LN994477 | 2 | 87 | 1089 | novel | - | - |
| *Mafa-B\*041:02* | KP307858 | 4 | 28 | 1089 | extant | 1089 | KP307858, KP307874, KP307882, LC053850 |
| *Mafa-B\*041:03* | LN851987 | 1 | 9 | 1089 | extension | 1071 | KP307846 |
| *Mafa-B\*043:01* | LN994478 | 2 | 33 | 1089 | extension | 646 | AB569230 |
| *Mafa-B\*043:02* | LN994479 | 1 | 15 | 1089 | novel | - | - |
| *Mafa-B\*044:01:01* | KP307903 | 10 | 113 | 1080 | extant | 1080 | FM212817, HM235712, KP307903 |
| *Mafa-B\*044:01:03* | LN994480 | 1 | 43 | 1080 | novel | - | - |
| *Mafa-B\*044:04* | HM581964 | 1 | 47 | 1080 | extant | 1080 | HM581964 |
| *Mafa-B\*045:07* | LN994481 | 2 | 21 | 1089 | novel | - | - |
| *Mafa-B\*047:03* | LN994482 | 1 | 53 | 1089 | novel | - | - |
| *Mafa-B\*048:06* | LN851988 | 1 | 54 | 1089 | novel | - | - |
| *Mafa-B\*050:02:02* | LN994483 | 1 | 15 | 1080 | novel | - | - |
| *Mafa-B\*050:05* | LN851989 | 5 | 55 | 1080 | extension | 546 | HM161343, HQ131714 |
| *Mafa-B\*050:08* | LN851990 | 2 | 10 | 1080 | extension | 1003 | LC043337 |
| *Mafa-B\*056:01* | KP307904 | 10 | 717 | 1089 | extant | 1089 | AY958131, HQ131699, KP307904 |
| *Mafa-B\*056:04* | KP307848 | 1 | 29 | 1089 | extant | 1089 | KP307848 |
| *Mafa-B\*056:05:01* | LN994484 | 1 | 52 | 1089 | extension | 1056 | HM235703, HQ131733, KP307905 |
| *Mafa-B\*056:05:02* | LN994485 | 1 | 54 | 1089 | novel | - | - |
| *Mafa-B\*060:01* | EU203692 | 4 | 6 | 1080 | extant | 1080 | EU203692 |
| *Mafa-B\*060:04* | HM581968 | 5 | 12 | 1080 | extant | 1080 | AB569226, HM581968, HQ131693 |
| *Mafa-B\*060:12* | LN851991 | 1 | 3 | 1080 | novel | - | - |
| *Mafa-B\*060:13* | LN851992 | 12 | 57 | 1080 | novel | - | - |
| *Mafa-B\*060:14* | LN994487 | 1 | 5 | 1080 | novel | - | - |
| *Mafa-B\*060:16* | LN994486 | 1 | 8 | 1080 | novel | - | - |
| *Mafa-B\*061:03:01* | KP307851 | 2 | 48 | 1080 | extant | 1080 | KP307851 |
| *Mafa-B\*061:03:02* | LN851993 | 1 | 11 | 1080 | novel | - | - |
| *Mafa-B\*061:04:01* | KP307861 | 6 | 270 | 1080 | extant | 1080 | KP307861, KP307843 |
| *Mafa-B\*061:04:02* | LN851994 | 2 | 77 | 1080 | novel | - | - |
| *Mafa-B\*061:05* | LN994488 | 2 | 5 | 1089 | novel | - | - |
| *Mafa-B\*064:05* | LN994489 | 1 | 4 | 1077 | novel | - | - |
| *Mafa-B\*065:02* | LN851995 | 3 | 28 | 1089 | extension | 822 | HM161351, HQ113986, HQ131727 |
| *Mafa-B\*065:04* | LN851996 | 5 | 70 | 1089 | novel | - | - |
| *Mafa-B\*065:05* | LN994490 | 1 | 40 | 1089 | novel | - | - |
| *Mafa-B\*066:01* | KP307849 | 1 | 16 | 1089 | extant | 1089 | KP307849 |
| *Mafa-B\*067:01* | LN994491 | 1 | 47 | 1089 | novel | - | - |
| *Mafa-B\*067:02* | LN994492 | 1 | 31 | 1089 | novel | - | - |
| *Mafa-B\*068:02* | FM212839 | 4 | 79 | 1089 | extant | 1089 | FM212839 |
| *Mafa-B\*068:04* | LN851997 | 7 | 110 | 1089 | extension | 641, 1056 | AB569236, KP307906 |
| *Mafa-B\*068:06:01* | KP307865 | 6 | 120 | 1089 | extant | 1089 | KP307865 |
| *Mafa-B\*068:06:02* | LN994493 | 1 | 23 | 1089 | novel | - | - |
| *Mafa-B\*068:07:01* | KP307850 | 1 | 20 | 1089 | extant | 1089 | KP307850 |
| *Mafa-B\*068:07:02* | LN994494 | 2 | 57 | 1089 | novel | - | - |
| *Mafa-B\*068:10* | LN851998 | 1 | 5 | 1089 | novel | - | - |
| *Mafa-B\*069:03* | LN852001 | 3 | 144 | 1089 | extension | 544 | HQ131732 |
| *Mafa-B\*069:04* | LN851999 | 2 | 113 | 1089 | extension | 822 | HM161434, HQ114002, HQ131710 |
| *Mafa-B\*069:05* | LN852000 | 2 | 152 | 1089 | novel | - | - |
| *Mafa-B\*069:06* | LN994495 | 1 | 90 | 1089 | novel | - | - |
| *Mafa-B\*069:07* | LN994496 | 1 | 28 | 1089 | extension | 822 | HM161424 |
| *Mafa-B\*070:04* | LN852002 | 2 | 10 | 1080 | extension | 1070 | KP307842 |
| *Mafa-B\*072:02* | HM581961 | 1 | 3 | 1089 | extant | 1089 | HM581961 |
| *Mafa-B\*072:04* | LN852003 | 2 | 5 | 1089 | novel | - | - |
| *Mafa-B\*073:02* | KP307876 | 2 | 82 | 1089 | extant | 1089 | HM161413, KP307876 |
| *Mafa-B\*073:03* | LN994497 | 1 | 18 | 1089 | novel | - | - |
| *Mafa-B\*074:02* | LN852004 | 7 | 132 | 1080 | extension | 646 | AB569228 |
| *Mafa-B\*075:01* | EU203718 | 1 | 73 | 1080 | extant | 1080 | AY958150, EU203718, GU063741 |
| *Mafa-B\*077:01* | FM212843 | 1 | 88 | 1089 | extant | 1089 | FM212843 |
| *Mafa-B\*077:02* | LN852005 | 1 | 59 | 1089 | novel | - | - |
| *Mafa-B\*079:01:01* | AM943362 | 1 | 1 | 1080 | extant | 1080 | AM943362 |
| *Mafa-B\*079:02:01* | LN994499 | 5 | 9 | 1080 | extension | 822 | HM236276 |
| *Mafa-B\*081:01* | LN852006 | 2 | 31 | 1086 | extension | 822 | AB569225, HM161374, HQ131721 |
| *Mafa-B\*082:03* | LN994500 | 6 | 20 | 1116 | novel | - | - |
| *Mafa-B\*083:02* | KP307907 | 1 | 151 | 1089 | extant | 1089 | HM236289, KP307907 |
| *Mafa-B\*083:04* | LN994501 | 1 | 16 | 1089 | novel | - | - |
| *Mafa-B\*085:01* | KP307852 | 7 | 214 | 1089 | extant | 1089 | AY958124, HM161375, HQ113995, KP307852, LC043363 |
| *Mafa-B\*088:04* | LN852007 | 8 | 13 | 1089 | novel | - | - |
| *Mafa-B\*088:05* | LN994502 | 6 | 12 | 1089 | novel | - | - |
| *Mafa-B\*089:01:02* | EU392125 | 6 | 18 | 1116 | extant | 1116 | EU392125, FJ178818, HM161360, HQ113989, JF729417, LC043349 |
| *Mafa-B\*091:03* | LN994503 | 1 | 70 | 1089 | extension | 822 | HM161370 |
| *Mafa-B\*093:02* | LN852008 | 5 | 242 | 1089 | extension | 822 | HM161359, HQ113988, HQ114006, HQ131734 |
| *Mafa-B\*095:03* | LN994504 | 1 | 12 | 1089 | novel | - | - |
| *Mafa-B\*095:04* | LN994505 | 1 | 173 | 1089 | novel | - | - |
| *Mafa-B\*098:03:02* | LN994508 | 3 | 5 | 1089 | novel | - | - |
| *Mafa-B\*098:04* | LN994506 | 3 | 6 | 1089 | extension | 981 | HM581969 |
| *Mafa-B\*098:12* | LN994507 | 6 | 15 | 1089 | novel | - | - |
| *Mafa-B\*101:02* | LN994509 | 2 | 164 | 1089 | extension | 819 | HQ113999, HQ131702 |
| *Mafa-B\*103:01:02* | LN994510 | 1 | 6 | 1080 | novel | - | - |
| *Mafa-B\*104:01:02* | FM212841 | 6 | 698 | 1089 | extant | 1089 | FM212841, HQ131684, HQ131774 |
| *Mafa-B\*105:01* | LN852009 | 2 | 80 | 1089 | extension | 819 | HQ114013, HQ131692 |
| *Mafa-B\*110:01:01* | LN852010 | 4 | 162 | 1089 | extension | 822 | HM161339, HM161378, HQ131725 |
| *Mafa-B\*110:01:02* | LN994511 | 1 | 13 | 1089 | novel | - | - |
| *Mafa-B\*117:03* | LN852011 | 2 | 26 | 1080 | novel | - | - |
| *Mafa-B\*117:04* | LN994512 | 1 | 18 | 1080 | novel | - | - |
| *Mafa-B\*124:01:01* | EU203714 | 1 | 27 | 1080 | extant | 1080 | EU203714 |
| *Mafa-B\*124:01:03* | KP307862 | 6 | 105 | 1080 | extant | 1080 | KP307862, KP307847 |
| *Mafa-B\*124:03* | LN852012 | 2 | 9 | 1080 | novel | - | - |
| *Mafa-B\*124:04* | LN994513 | 1 | 6 | 1080 | novel | - | - |
| *Mafa-B\*137:03* | EU203723 | 6 | 138 | 1089 | extant | 1089 | EU203723, EU392117, HM235700, HQ131688, HQ131740, JF729416, LC043364 |
| *Mafa-B\*137:05* | LN852013 | 1 | 12 | 1089 | novel | - | - |
| *Mafa-B\*137:06* | LN994515 | 2 | 32 | 1089 | novel | - | - |
| *Mafa-B\*137:07* | LN994514 | 1 | 4 | 1089 | novel | - | - |
| *Mafa-B\*137:08* | LN994516 | 1 | 11 | 1089 | novel | - | - |
| *Mafa-B\*139:02* | FM212840 | 2 | 19 | 1086 | extant | 1086 | FM212840 |
| *Mafa-B\*140:01:02* | LN994517 | 1 | 27 | 1089 | novel | - | - |
| *Mafa-B\*144:01* | LN994518 | 2 | 38 | 1080 | extension | 1042 | AY958122, HQ113983 |
| *Mafa-B\*145:01* | LN852014 | 3 | 198 | 1089 | extension | 1042 | AY958123, HM235702, HQ113984, HQ131728 |
| *Mafa-B\*146:01:01* | LN852015 | 1 | 24 | 1089 | extension | 1042 | AY958125 |
| *Mafa-B\*148:01:01* | LN994519 | 1 | 22 | 1089 | novel | - | - |
| *Mafa-B\*149:01* | EU606047 | 2 | 4 | 1089 | extant | 1089 | EU606047 |
| *Mafa-B\*149:02* | LN852016 | 2 | 10 | 1089 | extension | 822 | HG977501, HQ110973 |
| *Mafa-B\*150:02* | LN994520 | 1 | 96 | 1089 | extension | 1056 | FM212814 |
| *Mafa-B\*155:01* | EU203684 | 1 | 45 | 1080 | extant | 1080 | EU203684, HM235713 |
| *Mafa-B\*156:01* | EU203694 | 2 | 57 | 1089 | extant | 1089 | EU203694, FJ178816 |
| *Mafa-B\*161:05* | LN852017 | 2 | 57 | 1089 | novel | - | - |
| *Mafa-B\*164:02* | GQ153333 | 1 | 49 | 1089 | extant | 1089 | GQ153333, LC043371 |
| *Mafa-B\*181:03* | LN994521 | 1 | 41 | 1089 | novel | - | - |
| *Mafa-B\*184:01:01* | LN994522 | 2 | 162 | 1089 | extension | 822 | JN032108 |
| *Mafa-B\*184:01:02* | LN994523 | 2 | 100 | 1089 | novel | - | - |
| *Mafa-B\*203:01* | LN994498 | 1 | 81 | 1089 | novel | - | - |
| *Mafa-I\*01:01:02* | LN994530 | 1 | 9 | 1089 | novel | - | - |
| *Mafa-I\*01:09* | AB195465 | 1 | 2 | 1089 | extant | 1089 | AB195465 |
| *Mafa-I\*01:11:02* | LN852020 | 14 | 46 | 1089 | novel | - | - |
| *Mafa-I\*01:14:02* | LN852021 | 4 | 5 | 1089 | novel | - | - |
| *Mafa-I\*01:18:01* | LN994524 | 8 | 34 | 1089 | extension | 1056 | FM246497 |
| *Mafa-I\*01:22* | HM581970 | 1 | 2 | 1089 | extant | 1089 | AF161864, HM581970 |
| *Mafa-I\*01:29* | LN852018 | 2 | 6 | 1089 | novel | - | - |
| *Mafa-I\*01:30* | LN852019 | 4 | 4 | 1089 | novel | - | - |
| *Mafa-I\*01:31* | LN994525 | 4 | 6 | 1089 | novel | - | - |
| *Mafa-I\*01:32* | LN994526 | 3 | 6 | 1089 | novel | - | - |
| *Mafa-I\*01:33* | LN994527 | 4 | 8 | 1089 | novel | - | - |
| *Mafa-I\*01:34* | LN994528 | 1 | 6 | 1089 | novel | - | - |
| *Mafa-I\*01:35* | LN994529 | 1 | 6 | 1089 | novel | - | - |

Summary of 313 full-length MHC class I transcripts identified in 72 cynomolgus macaques from Chinese breeding facilities

All PCR products were purified using the AMPure XP PCR purification kit (Agencourt Bioscience Corporation, Beverly, MA, USA) and quantified using the Quant-iT dsDNA HS Assay kit and a Qubit fluorometer (Invitrogen, Carlsbad, CA, USA), following the manufacturer’s protocols.

### Sequencing

The MHC class I exon 2 genotyping amplicons were sequenced on an Illumina MiSeq instrument (San Diego, CA, USA), and the full-length MHC class I transcripts were sequenced on a PacBio RS II instrument (Menlo Park, CA, USA) following the respective manufacturer’s protocols. Additional sequencing details are provided in Supplemental File 1, and an overview of the PacBio sequencing process is shown in Supplemental Fig. 3.

**Fig. 3.**
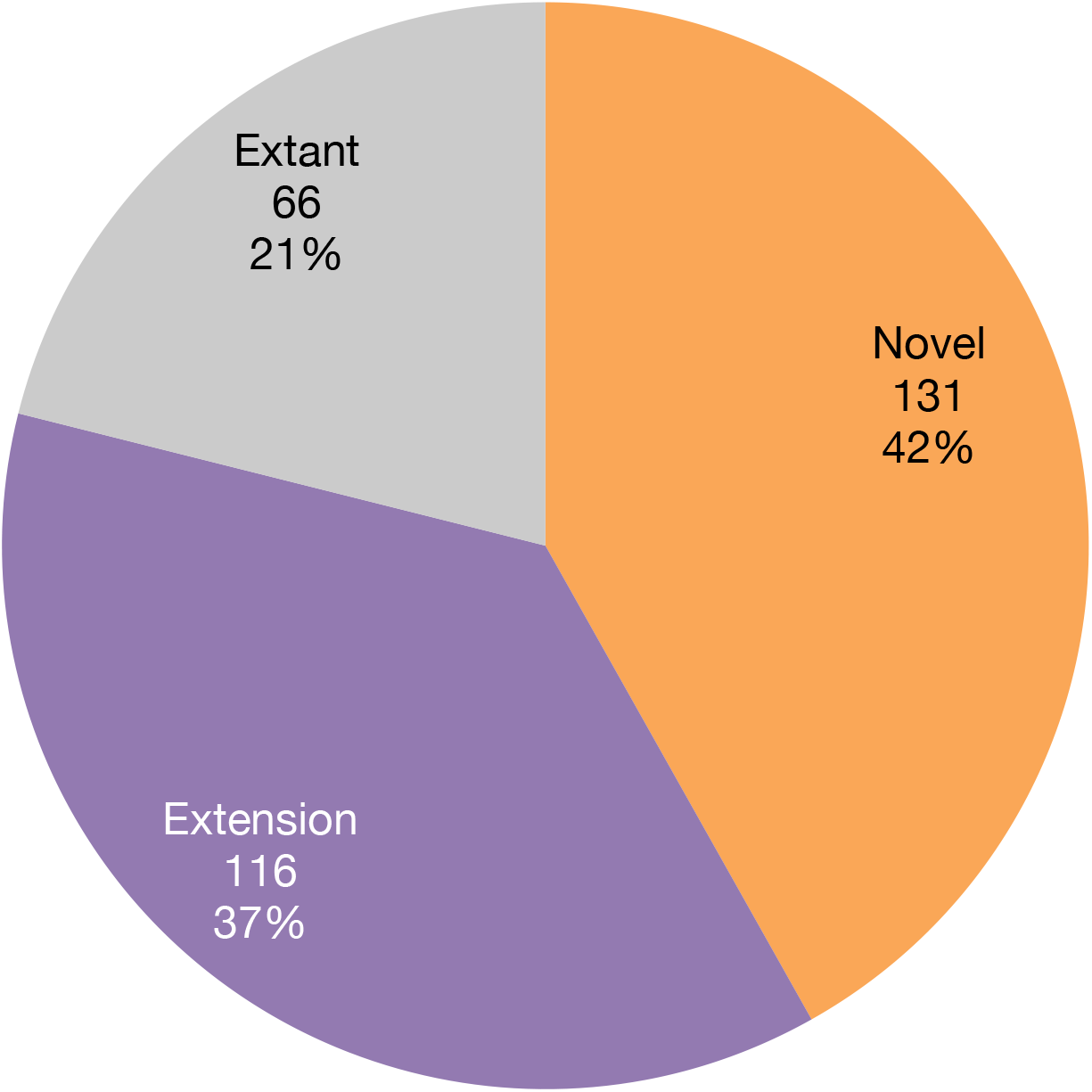
Evaluation of PacBio full-length transcripts. A breakdown of the 313 total identified *Mafa-A*, *Mafa-B*, and *Mafa-I* full-length transcripts

### Data analysis

Processing of the MHC class I exon 2 genotyping amplicon was performed as described previously for MHC class II amplicons (Karl et al. 2014) using a custom-built command line pipeline to merge reads, trim primers, and compare MiSeq reads against a database of known cynomolgus macaque transcripts. *Mafa-A* and *Mafa-B* lineage-level haplotypes were determined for each of the 100 samples essentially as previously described (Heimbruch et al. 2015; Karl et al. 2013), first examining the sire/dam/progeny trio sets, then the parent/offspring duos, to look for sets of co-inherited sequences. Haplotypes for the unrelated animals in this cohort were inferred based on similarity to those observed in the related sets of animals.

A custom semi-automated pipeline was also developed for analysis of the PacBio MHC class I full-length transcripts; an overview of this pipeline is shown in Supplemental Fig. 4. Full details of the pipeline are available in Supplemental File 1, and the scripts and tools used are publicly available (https://bitbucket.org/dholab/mhc-long-amplicon-allele-discovery-in-mafa-from-china). In brief, the PacBio reads of insert (the consensus sequence generated from all passes around the circular SMRTbell template) were processed to eliminate indel sequences, short reads, and chimeric sequences using the SMRT Analysis v2.3 Reads of Insert protocol. Processed reads perfectly matching known full-length cynomolgus macaque transcripts were removed prior to clustering by using bbmap (version 34) to map reads to known full-length allele sequences. Reads not identical to known allele sequences were clustered using USEARCH–cluster_fast (version v8.0.1517_i86osx32) and requiring 100% identity within clusters. Clusters supported by three or more identical full-length reads were then binned as either extensions of known partial cynomolgus macaque sequences or putative novel transcripts, and a genotyping table with read counts per sample was generated. All extension and putative novel sequences were manually validated using Geneious Pro v9.0 (Biomatters Limited, Auckland, New Zealand) to perform BLAST alignments and create BAM alignment files of the reads of insert against the validated transcripts. Validated sequences were submitted to GenBank and the Immuno Polymorphism Database for the Major Histocompatibility Complex genes of Non-Human Primates (IPD-MHC NHP) (http://www.ebi.ac.uk/ipd/mhc/nhp/index.html) (de Groot et al. 2012; Robinson et al. 2013) for official nomenclature.

### Data availability

Primer sequences are listed in Supplemental Fig. 1 and 2. Lineage-level *Mafa-A* and *Mafa-B* haplotypes assigned to each animal from the MiSeq exon 2 data are available in Supplemental Table 1, and full MiSeq read counts per transcript per animal are available in Supplemental Fig. 5. Full resolution *Mafa-A* and *Mafa-B* haplotypes assigned to each of the 72 animals examined by PacBio sequencing are available in Supplemental Table 2, and full PacBio read counts per transcript per animal are available in Supplemental Fig. 6. Novel and extension full-length sequences as listed in Table 1 are available from GenBank (LN851916-LN852021, LN994391-LN994530, LN998188, KY047492).

## RESULTS AND DISCUSSION

### Lineage-level haplotype analysis

In human and macaque MHC class I nomenclature, closely related alleles originating from the same gene are assigned to the same lineage group, signified by the 2-3 digits between the asterisk and the first colon of MHC class I sequence names. Since the Illumina MiSeq exon 2 genotyping amplicon is focused on a small segment of the MHC class I molecule, it is generally only sufficient for determining the lineage groups of any detected MHC class I sequences (considered ‘three-digit’ resolution in human HLA genotyping and referred to as ‘lineage-level’ resolution here). However, it provides a rapid, high-throughput methodology for haplotype analysis, especially within related animals. Using the parent/progeny trio and duo sets within these two cohorts, we described haplotypes of *Mafa-A* and *Mafa-B* transcripts inherited together from parents to offspring. Two representative trio sets with parents and their progeny are illustrated in Fig. 1a. Any transcripts observed as shared between parent and offspring were assigned to the inherited haplotype; any remaining transcripts observed within a parent, or in an offspring with just one representative parent, were assigned to the alternate haplotype. For animals without a direct relative in the cohort, haplotypes were inferred based on identity to the defined inherited haplotypes. We focused on the ‘major’ transcripts with high levels of steady state RNA for haplotype definitions. Abbreviated haplotype names were assigned based on presence of a major ‘diagnostic’ sequence that was typically the most abundant transcript on the haplotype, i.e., B013 haplotypes express the diagnostic major transcript *Mafa-B*013*. Different combinations of major allele lineages inherited with each diagnostic transcript were denoted with lowercase letters following the haplotype name, e.g., *Mafa-B*013* inherited with *Mafa-B*007* was designated B013a and *Mafa-B*013* inherited with *Mafa-B*137* was designated B013d. The full list of *Mafa-A* and *Mafa-B* haplotypes defined in these cohorts of cynomolgus macaques from Chinese breeding facilities is given in Supplemental Table 1.

A total of 52 distinct *Mafa-A* haplotypes and 74 unique *Mafa-B* haplotypes were identified in the 100 animals (200 chromosomes) evaluated in this cohort (Supplemental Table 1). A summary of the lineage-level haplotypes observed in each animal is shown in Fig. 2, with haplotypes directly inherited from parent to offspring denoted with solid outlined boxes. Full MiSeq haplotyping results, including reads supporting each transcript call per sample, are also given in Supplemental Fig. 5. While overall the large number of related animals in the cohort skews haplotype frequencies, it should be noted that four *Mafa-A* haplotypes (A006, A022a, A027, and A065) and three *Mafa-B* haplotypes (B013d, B028d, and B039e) are each observed in ten or more individuals. There are 1-2 major alleles expressed on each *Mafa-A* haplotype, accompanied by 0-5 minor alleles. Each *Mafa-B* haplotype is associated with 1-7 expressed major transcripts and between 0-15 minor transcripts.

Though many studies have examined the lists of MHC class I transcripts expressed in cynomolgus macaques of multiple geographic origins, few have studied the haplotype structures in those animals (Budde et al. 2010; Campbell et al. 2008; Otting et al. 2009, 2012; Saito et al. 2012; Shiina et al. 2015). The best-characterized population of cynomolgus macaques is from the island of Mauritius, where a limited number of founding animals were deposited on the island in the 1500s. Since the entire current population arose from a small number of ancestors, the diversity of their MHC region (and their entire genome) is limited to five lineage-level *Mafa-A* haplotypes (A031, A032a, A033, A060, A063) and seven lineage-level *Mafa-B* haplotypes (B019a, B045a, B075a, B104a, B147, B151a, B164a) (Budde et al. 2010; Wiseman et al. 2013). The diversity observed in these cohorts of 100 individuals from two Chinese breeding facilities represents a ten-fold increase in diversity at each locus compared to Mauritian cynomolgus macaques. While both populations of cynomolgus macaques are popular for biomedical research, this difference in overall diversity suggests that investigators should carefully consider their objectives when choosing a source for their animals. For protocols where MHC may be a confounding factor, like SIV and transplantation research, the restricted diversity of the Mauritian cynomolgus macaque population is particularly attractive. When testing vaccines or other drug therapies for potential negative side effects however, an extremely diverse source like cynomolgus macaques from Chinese breeding facilities may be advantageous. The overall diversity observed in this cohort also suggests that researchers should have their animals haplotyped if the MHC is a possible modifying factor, since this study revealed a large number of haplotypes despite the fact that multiple related animals were evaluated.

This study likely represents a snapshot of the total MHC diversity available within cynomolgus macaques from Chinese facilities, largely dependent on the country of origin of their breeding animals. Of the 52 total *Mafa-A* haplotypes observed here, nine (17%) were observed exclusively in animals of reported Vietnamese heritage, 17 (33%) were of reported Cambodian-origin only, and 4 (8%) were observed in the small set of reported hybrid Cambodian/Indonesian-origin animals (Supplemental Table 1). Only 22 (42%) *Mafa-A* haplotypes were observed in samples from multiple origins. The results for the 74 *Mafa-B* haplotypes are even more striking with only 21 (28%) observed in samples from multiple origins. Among the remaining *Mafa-B* haplotypes, 14 (19%) were exclusive to Vietnamese-origin animals, 26 (35%) exclusive to Cambodian-origin, and 13 (18%) were observed only in the mixed Cambodian/Indonesian-origin individuals. No previous MHC haplotype studies have been performed in either Vietnamese–or Cambodian-origin cynomolgus macaques, but limited studies have characterized haplotypes in Filipino, Indonesian, and Malaysian cynomolgus macaques (Campbell et al. 2008; Otting et al. 2009, 2012; Saito et al. 2012; Shiina et al. 2015). The cohorts studied here appear more MHC diverse than those sampled from the previously described insular and extremely southern continental Asian populations (it is unclear from precisely where the Malaysian cohort was initially derived).

### Full-length allele discovery

In this study we employed a novel method for next-generation sequencing of full-length MHC class I alleles, using the PacBio RS II SMRT long read sequencing instrument. In our previous study with P4-C2 PacBio sequencing (Westbrook et al. 2015), data analysis relied on creation of contigs at 99% identity (largely to account for insertion/deletion artifacts) to define transcripts. Since this initial study, advances in PacBio sequencing with P6-C4 reagents and analysis software have eliminated the need to include less than 100% identical reads for identification of novel sequences. From the full 100 animal cohort, we selected 72 total samples across both breeding centers for PacBio sequencing based on lineage-level haplotype diversity. Table 1 presents a complete list of all 313 full-length MHC class I transcripts that we detected in these 72 individuals; these include 126 *Mafa-A*, 174 *Mafa-B*, and 13 *Mafa-I* transcripts. The complete PacBio sequencing results for each animal including numbers of identical sequencing reads supporting each transcript call are shown in Supplemental Fig. 6. Overall, nearly 80% of the transcripts identified in this study added new information to the existing *Mafa* MHC class I allele databases – 131 (42%) of the sequences were completely novel to the database, and 116 (37%) extended short but previously named database sequences to cover full-length open reading frames (Fig. 3).

This study improved the extant cynomolgus macaque allele databases in both total number of sequences and, more importantly, in the percentage of known transcripts that have been unambiguously described over the full coding region. The November 2015 update of IPD-MHC NHP contained a total of 899 *Mafa-A*, *Mafa-B*, and *Mafa-I* sequences. Of these, only 386 (43%) were full-length from start codon to stop codon. This study adds an additional 131 novel transcripts to the database, bringing the total to 1,030 named MHC class I sequences in the cynomolgus macaque database. With the additional 116 extant sequences extended to full length, the database of *Mafa-A*, *Mafa-B*, and *Mafa-I* transcripts following this study now contains 633 full-length sequences (61% of the 1,030 total transcripts).

We also examined distinct, full resolution haplotypes in the samples that were PacBio sequenced to compare to the previously determined lineage-level haplotypes from the full cohort. Fig. 1b shows the PacBio haplotype results for two representative trio sets of samples, and Supplemental Table 2 provides the complete list of full resolution haplotypes observed in the cohort. Haplotype names were retained from the lineage-level haplotype results, but sub-haplotypes (indicated with Roman numerals in the “Sub-Haplo” column of Supplemental Table 2) were assigned for any distinctions in specific transcripts. For instance, the lineage-level haplotype A001 has four (i-iv) distinct sub-haplotypes in the PacBio data, as they all vary from each other at the transcript level (A001i – A001iv carry *Mafa-A1*001:01:02*, *Mafa-A1*001:01:03*, *Mafa-A1*001:02:01*, and *Mafa-A1*001:05*, respectively). For the PacBio haplotypes, both major and minor specific transcripts were considered when assigning sub-haplotypes. As expected, results were concurrent with the lineage-level haplotypes overall, but the full-length allelic resolution provided by PacBio sequencing resulted in a significant expansion of the number of both *Mafa-A* and *Mafa-B* haplotypes. In total we identified 92 and 101 distinct full-resolution *Mafa-A* and *Mafa-B* haplotypes, respectively, in the 72 PacBio-sequenced samples. As with the lineage-level haplotypes, more than 90% of the *Mafa-A* and *Mafa-B* PacBio haplotypes were only observed in individuals from a single origin– 26 *Mafa-A* and 29 *Mafa-B* haplotypes were distinct to Vietnamese-origin samples, 43 *Mafa-A* and 46 *Mafa-B* haplotypes were only observed in Cambodian-origin samples, and 15 *Mafa-A* and 15 *Mafa-B* haplotypes were distinct to the hybrid Cambodian/Indonesian-origin samples (Supplemental Table 2).

It is unclear how many of these allelic distinctions make a functional biological difference in regards to immune responses. Although such confirmatory studies are beyond the scope of this report, we predict that specific nonsynonymous nucleotide variants between two closely related alleles may result in significant biological differences, particularly when the nucleotide differences alter the resulting amino acid within the peptide binding pocket. Therefore, full resolution genotyping provided by PacBio sequencing is particularly attractive for researchers looking for linkage between a specific immune response and specific MHC class I genotypes.

Overall, this study describes some of the first MHC class I *Mafa-A* and *Mafa-B* haplotypes in cynomolgus macaques from Chinese breeding facilities, including samples of reported Vietnamese, Cambodian, and mixed Cambodian/Indonesian origins. It also describes an improved PacBio long read sequencing approach to characterize full-length MHC class I transcripts without contig assembly, using only the highest quality reads. The transcripts identified here both increase the total number of known *Mafa-A*, *Mafa-B*, and *Mafa-I* sequences, and significantly improve the percentage of full-length transcripts in the nonhuman primate IPD database. Finally, the general PacBio-based method described here for long-amplicon sequencing can also be applied to other complex immune loci of macaques such as MHC class II, killer immunoglobulin-like receptors, and Fc gamma receptors, as well as those of other model organisms.

## ACKNOWLEDGMENTS

The authors thank Catherine Westbrook, Suzanne Mate, and Galina Koroleva at the U.S. Army Medical Research Institute of Infectious Diseases, and Katherine Munson and the staff of the University of Washington PacBio Sequencing Services core for performing PacBio RS II sequencing. We also thank Annemiek J.M. de Vos-Rouweler for RNA isolations and preparation of cDNAs used in this study. Finally, we thank Nel Otting, Natasja de Groot and the IMGT Non-human Primate Nomenclature Committee for obtaining accession numbers and providing official *Mafa* transcript nomenclature designations for the sequences described here.

This research was supported by contracts HHSN272201600007C and HHSN272201100013C from the National Institute of Allergy and Infectious Diseases of the National Institutes of Health, and was conducted at a facility constructed with support from the Research Facilities Improvement Program (RR15459-01, RR020141-01).

